# An Artificial Intelligence Method for Phenotyping of OCT Scans Using Unsupervised and Self-supervised Deep Learning

**DOI:** 10.1101/2023.10.20.563205

**Authors:** Saber Kazeminasab, Sayuri Sekimitsu, Mojtaba Fazli, Mohammad Eslami, Min Shi, Yu Tian, Yan Luo, Mengyu Wang, Tobias Elze, Nazlee Zebardast

**Author notes:** Correspondence: Nazlee Zebardast MD MS.

## Abstract

Artificial intelligence (AI) has been increasingly used to analyze optical coherence tomography (OCT) images to better understand physiology and genetic architecture of ophthalmic diseases. However, to date, research has been limited by the inability to transfer OCT phenotypes from one dataset to another. In this work, we propose a new AI method for phenotyping and clustering of OCT-derived retinal layer thicknesses using unsupervised and self-supervised methods in a large clinical dataset using glaucoma as a model disease and subsequently transfer our phenotypes to a large biobank. The model includes a deep learning model, manifold learning, and a Gaussian mixture model. We also propose a correlation analysis for the performance evaluation of our model based on Pearson correlation coefficients. Our model was able to identify clinically meaningful OCT phenotypes and successfully transfer phenotypes from one dataset to another. Overall, our results will contribute to stronger research methodologies for future research in OCT imaging biomarkers, augment testing of OCT phenotypes in multiple datasets, and ultimately improve our understanding of pathophysiology and genetic architecture of ocular diseases.

## Introduction

Advances in artificial intelligence (AI), machine learning (ML), and deep learning (DL) have allowed for more precise analysis of medical imaging with the goals of better understanding disease pathogenesis and predicting disease burden. In particular, image phenotyping has enabled the extraction of complex patterns and subtle characteristics from medical imaging data, thereby unlocking novel insights into disease mechanisms and paving the way towards personalized diagnostic and therapeutic approaches. The majority of the medical images are unlabeled as the labeling task is expensive and time-consuming. Hence, methods like unsupervised learning or self-supervised learning are used for pattern recognition. Unsupervised learning employs machine learning algorithms to identify features of unlabeled datasets. These datasets are analyzed and clustered without the necessity of explicit labels or annotations. This allows the model to autonomously recognize and learn patterns within the input data. Through these methods, similarities and differences between data features can be discerned directly from the data itself. Techniques such as non-parametric instance discrimination (1), DeepCluster (2), unsupervised deep embedding (3), autoencoders (4), and deep adaptive image clustering (5) serve as examples of unsupervised learning for clustering tasks. Selfsupervised learning extracts the feature space by designing a proxy task derived from the data itself, which is a noted limitation of this approach (6–10). This challenge, mainly the design of the proxy task in self-supervised learning, is predominantly addressed by contrastive learning (11). The objective in contrastive learning is to optimize a function for similar pairs in contrast to dissimilar pairs (12–21). The use of selfsupervised learning on medical images is well studied in the literature (10, 22–26). Prior work has demonstrated the utility of image phenotyping in various ophthalmic applications such as glaucoma assessment (27), retinal disease diagnosis (28), genome-wide analysis study (GWAS) for vertical cup to disc ratio (VCDR) (29), as well as classification of agerelated macular degeneration (AMD), (30) and geographic atrophy (31). Technological advances in AI and their applications to medical imaging have been fueled by the advent of large-scale biobank studies with multimodal data (e.g., the UK Biobank) (32). However, to date, image analysis has largely been limited by the lack of data annotation by experts (33) and limited transferability from one dataset to another. These limitations have consequences in settings where datasets lack the information needed for specific studies. Improving the ability to transfer phenotypes from dataset to dataset may allow for more studies tying imaging phenotypes to genetic studies; for example, making it possible to apply a model trained on a large clinical dataset without genetic information to a population-based biobank study with limited clinical data but a robust genetic repository. Optical coherence tomography (OCT) is a non-invasive imaging technique which segments retinal layers, is used in the screening, diagnosis, and management of multiple ophthalmic disease including glaucoma. Glaucoma, a leading cause of irreversible vision loss and blindness worldwide, is a complex multifactorial disease with high degrees of structural and functional variation. Primary open-angle glaucoma (POAG), the most common subtype of glaucoma, is highly heritable (34). However, to date, POAG pathophysiology and genetic architecture remains incompletely explained. Clinicians often struggle with predicting which patients may progress to more severe disease. This is likely due significant structural and functional variation in glaucoma. While OCT has emerged as a valuable tool in clinical and research contexts, contributing significantly to our understanding of glaucoma (35), current image analysis methods do not capture disease heterogeneity. We propose that unsupervised and semi-supervised learning algorithms can used to identify OCT-based endophenotypes that reflect subtypes of glaucomatous optic neuropathy. The purpose of this study therefore is to use glaucoma as a model disease to describe a new AI method for phenotyping and clustering of OCT-derived retinal layer thicknesses using unsupervised and self-supervised methods. We also describe a new method for transferring and testing OCT phenotypes from one dataset to another. Our results will enhance future research in OCT imaging biomarkers, augment testing of OCT phenotypes in multiple datasets, and ultimately improve our understanding of pathophysiology and possibly genetic architecture of ocular diseases.

## Methods

### Massachusetts Eye and Ear and the UK Biobank

We used data from Massachusetts Eye and Ear Infirmary (MEE), a large tertiary-care center in the United States, and the UK Biobank (UKBB), a prospective cohort study of UK residents. The MEE dataset is a large collection of macular OCTs and associated clinical data from a diverse patient population with glaucoma. Individuals with glaucoma were identified by the International Classification of Diseases, Ninth or Tenth Revision (ICD 9/10) diagnosis code for glaucoma (ICD9: 365.x, ICD10: H40.x). The UKBB includes detailed genotypic and phenotypic information on over 500,000 participants from across the United Kingdom, including ophthalmic testing and OCT imaging on a subset of participants (32).

In the MEE dataset, we used data from subset of patient with Cirrus OCT scans of the macula (Zeiss, Inc, Germany). Three-dimensional macular volume scans (128 or 200 B-scans, with a configuration of 512 or 200 horizontal A-scans in a 6x6-mm raster pattern) were obtained. Images and OCT thickness layers were reconstructed using Zeiss’s automated boundary segmentations. This algorithm operates by analyzing the intensity and reflectance patterns of the incoming OCT scans. This algorithm uses complex image processing techniques, such as edge detection and pattern recognition, to identify and precisely delineate the boundaries between various retinal layers (36). In the UKBB, spectral-domain OCT scans of the macula were obtained on a subset of participants using Topcon 3D OCT 1000 Mk2 (Topcon, Inc, Japan). Three-dimensional macular volume scans were obtained (512 horizontal A-scans/B-scans; 128 B-scans in a 6x6-mm raster pattern). All OCT images were stored in .fda image files and the Topcon Advanced Boundary Segmentation (TABS) algorithm was used to automatically segment all scans. TABS uses dual-scale gradient information to allow for automated segmentation of the inner and outer retinal boundaries and retinal sublayers. For the MEE dataset, poor-quality images were identified. Missing pixels were imputed by the MissForest algorithm in Python (37). Also, the images with more than 5% missing values were excluded. The OCT scans in the UKBB have all pixels available.

### A. Unsupervised Artificial Intelligence Model

Our AI model is comprised of a DL model, manifold learning for dimensionality reduction, Gaussian Mixture Model (GMM) for clustering and defining components coefficients for each image, and a visualization tool that samples representative images for each cluster (**Figure 1**). The input for the models included (1) RNFL thickness maps and (2) GCC thickness maps. Unsupervised deep learning (DL) models developed in this work include a deep autoencoder and an encoder trained using unsupervised representation method. All models were trained on the MEE dataset and tested on both the MEE and UKBB datasets.

**Fig. 1.**
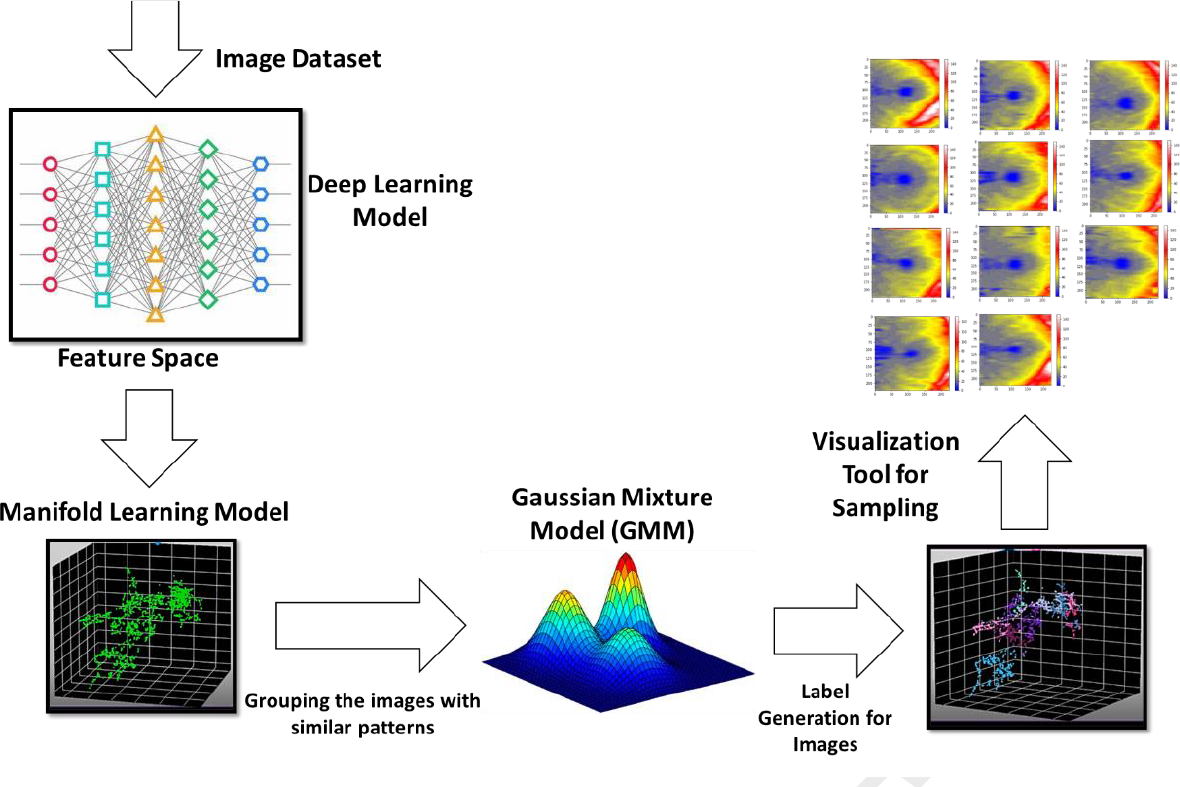
The proposed artificial intelligence (AI) model for macular OCT retinal layer thickness map phenotyping.

#### A.1 Deep Learning Models

We use two deep learning model in our study:

- Unsupervised Autoencoder based Visual Feature Extraction
- Encoder based on Self-Supervised Representation Learning

##### Unsupervised Autoencoder based Visual Feature Extraction

Since the scans do not have labels, we use unsupervised learning as one method for feature extraction. An autoencoder is a type of DL model used for It consists of an encoder and a decoder and works based on unsupervised learning. The encoder takes input data and maps it to a lower-dimensional latent space representation, while the decoder takes the latent space representation and reconstructs the original input. The autoencoder is trained by minimizing the difference between the input and the reconstructed output, using techniques such as backpropagation and stochastic gradient descent (38). **Figure 2** shows the schematic of the autoencoder we used in our work and **Supplemental Table S1** shows its parameters. The autoencoder model was implemented in the Keras framework.

**Fig. 2.**
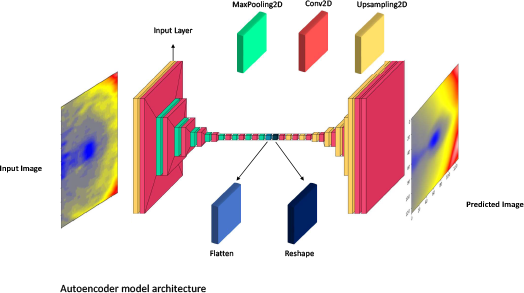
The autoencoder model for feature extraction of retina layers with unsupervised learning method.

##### Encoder based on Self-Supervised Representation Learning

Another way of feature extraction of the dataset is selfsupervised learning. We use momentum contrast (MOCO) (12) in which a feature model (e.g., an encoder) is trained based on a self-supervised learning method with contrastive learning as the proxy task. In this method, a dynamic dictionary for contrastive unsupervised learning is built using a queue and moving-averaged encoder. In MOCO, two random transformations of the same image are fed into two branches in which each branch includes an encoder and a projection head that is a two-layer perceptron. The encoder in the lower branch is only optimized during training and the encoder in the upper branch is considered a moving average of the encoder in the lower branch.

For training the encoder, let’s denote two transformations, *x*^′^ and *x*^″^ from the input image *x*. The outputs of the upper and lower branches are denoted as *u*^+^ = ℳ (ℱ (*x*^′^; *θ*′ℱ); *θ*′ℳ) and *u* = ℳ (*F* (*x*; *θ*ℱ); *θ*ℳ), respectively. *θ*′ℱ and *θ*′ℳ are the parameters of the feature model and projection head of the upper branch whereas *θ*ℱ and *θ*ℳ are the parameters for feature model and projection head of the lower branch. The loss function for this model is as follows:

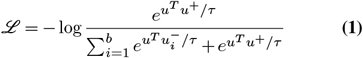

Where *b* is the batch size, and *u*^−^ is the output of lower branch for any image other than the image *x. τ* is the softmax temperature and is set to 0.2. The parameters of the models are updated in each iteration as *θ*^′^ℱ ← *µθ*^′^ℱ + (1 − *µ*)*θ*ℱ and *θ*′ ℳ← *µθ*′ℳ + (1 − *µ*)*θ*ℳ . *µ* is momentum coefficient and is set to 0.999. After training, the feature model and a projection head are accessible that transform the input image to a feature vector. S2 shows the parameters of the models and the hyperparameters for training based on the MOCO.

#### A.2 Manifold Learning for Dimensionality Reduction

Given that the features extracted from the previous DL models are high-dimensional, we used uniform manifold approximation and projection (UMAP) to project outputs from the previous model to a lower dimensional space (39, 40). The UMAP algorithm is a non-linear dimensionality reduction technique (41). It provides a unique representation of high-dimensional data, with underlying assumptions that the data is distributed uniformly on Riemannian manifold with a metric that is locally constant, and that the manifold is locally connected. Based on these assumptions, the manifold for the data is modeled with fuzzy topological structure in which the embedding (low dimensional representation) is constructed with the closest possible equivalent fuzzy topological structure (41). In our experiments, the latent space of 16,384 elements for the output of the autoencoder DL model and 128 elements for the encoder DL model were embedded into three elements for each data sample.

#### A.3 Number of Clusters Determination

As our dataset was unlabeled, we used the Bayesian information criterion (BIC) (42) and Akaike information criterion (AIC) for optimal selection of the number of phenotype clusters (43). BIC and AIC are commonly used statistical measures for model selection. Both BIC and AIC are based on the likelihood function of the model and include a penalty term to account for the number of parameters in the model. In general, models with lower AIC or BIC values are preferred over models with higher values, as they are considered to be more parsimonious. We calculated the AIC/BIC scores for the models with 2-20 clusters for each image type and DL model.

#### A.4 Gaussian Mixture Model (GMM) for Clustering

The Gaussian mixture model (GMM) is a statistical model that represents the probability distribution of a random variable as a weighted sum of Gaussian distributions (44). The probability density function (PDF) in one dimension for a Gaussian Distribution is expressed as:

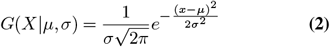

Where *µ* is the mean and *σ*^2^ is the variance of the distribution. Analogous to one-dimensional distribution, the PDF for a multivariate Gaussian distribution is given as:

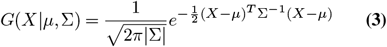

In Eq. 3, *µ* is d-dimensional vector indicating the mean of the distribution and is a *d × d* covariance matrix. Assume we have *K* clusters, we compute the *µ* and for each cluster. If *K* = 1, they are estimated by the maximum likelihood method. However, for *K >* 1, we have:

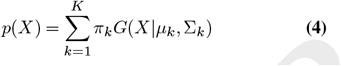

*π*_*k*_ is the mixing coefficient for the *k*^*th*^ distribution. More details about this method can be found in (45). In this work, we fit the GMM model on one embedding space and either predict the same embedding space or a different one. Also, we calculate the probability for each image in a particular cluster. The cluster label is assigned based on the maximum probability of the image in a cluster.

## Results

### Study Population

In the MEE dataset, we identified 18,985 images of 8,323 patients with a diagnosis code for glaucoma. Of the 8,323 patients, 43.81% were men with an average age of 61.63 +/15.98 years. The race/ethnicity distribution of the study population is as follows: 57.47% White, 16.11% Black, 7.27% Asian, 0.16% American Indian or Alaska Native, 0.11% Native Hawaiian or Other Pacific Islander, and 13.81% Hispanic.

In the UKBB dataset, we identified 86,115 images of 47,908 participants. Of the 47,908 participants, 45.8% were male with an average age of 56.39 +/8.08 years. The ancestry distribution of the study population is as follows: 91.3% European ancestry, 3.95% African ancestry, 3.47% South Asian ancestry, 0.83% East Asian ancestry, and 0.43% American ancestry.

### Autoencoder Model

The reconstruction error of trained autoencoders were computed (**Figure 3**), based on the comparison of each pixel of the original image and the reconstructed image. We calculated the error for each image and computed the mean error across all images. For MEE, the reconstruction error for RNFL and GCC was 18% and 6%, respectively. For UKBB, the reconstruction error for RNFL and GCC was 20% and 9%, respectively (**Figure 3**). The similar reconstruction errors for MEE and UKBB indicate that the autoencoder was able to successfully learn RNFL and GCC patterns identified in the MEE dataset and map them to the UKBB dataset.

**Fig. 3.**
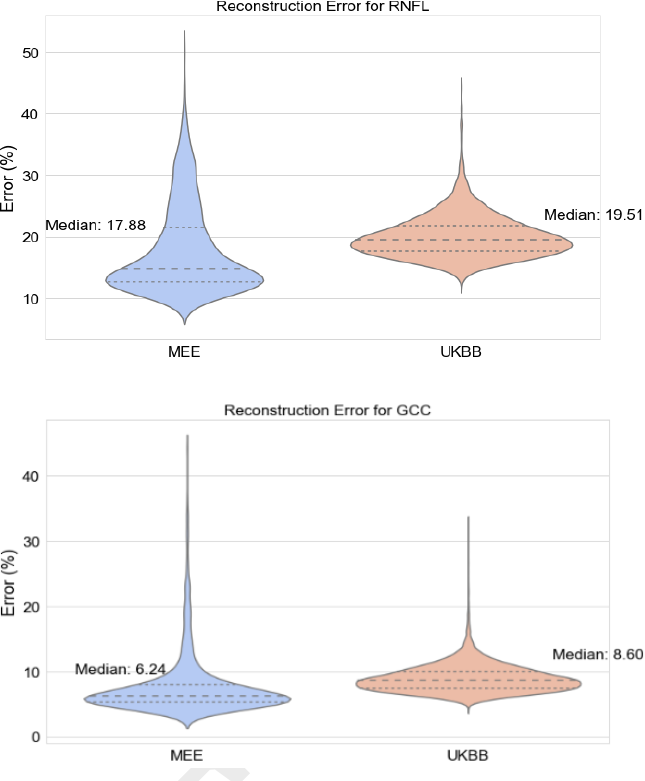
Reconstruction error for the autoencoders that are trained on GCC and RNFL images of Mass Eye and Ear (MEE) dataset.

We applied UMAP to the predictions of the autoencoder to reduce the dimensionality of the data samples. The parameters of the UMAP model are listed in **Supplemental Table S3**. AIC and BIC scores for embedding were calculated and three candidates for the optimal number of clusters are considered. The optimal number of clusters was found to be 11 for GCC and 9 for RNFL. This was selected based on a correlation analysis between images within and between clusters.

Different numbers of clusters were tested; the number of clusters with the highest correlation within images of a clusters compared to correlation of images between clusters were chosen (**Figure 4**). The applied GMM on the embeddings is shown in **Supplemental Figure S1**. Labels for the output of the GMM models were created. The labels are shown in the embedding space of MEE dataset for GCC and RNFL images in **Figure 5**. Representative sample images from each cluster are shown in **Figure 5**. The weight of the patterns for each dataset is shown in **Figure 6**. As seen in **Figure 6**, some patterns seen in MEE were not seen when the model was tested in the UKBB (RNFL – pattern 8, GCC – patterns 1, 5).

**Fig. 4.**
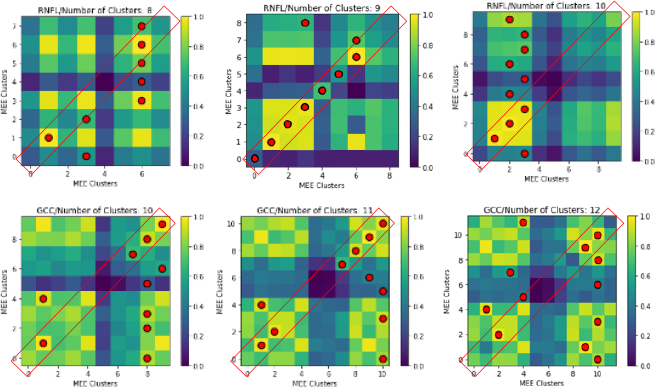
The correlation analysis between the images within a cluster and the images in different clusters for different numbers of clusters for the autoencoder model.

**Fig. 5.**
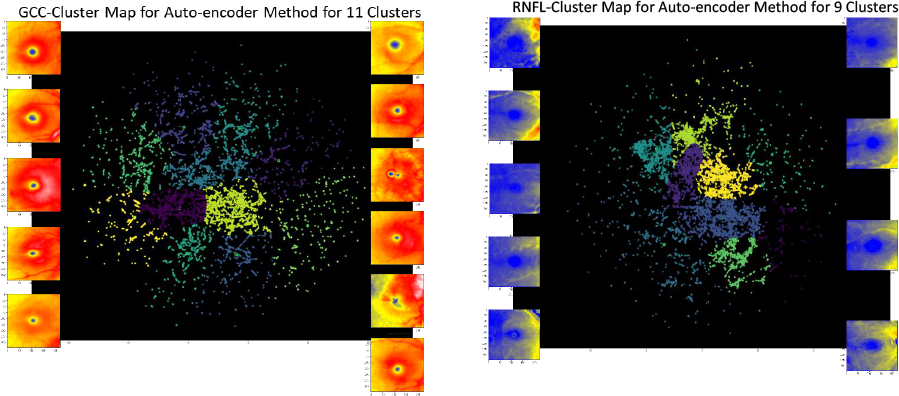
Labels of data points in the embedding space of GCC and RNFL images of autoencoder model.

**Fig. 6.**
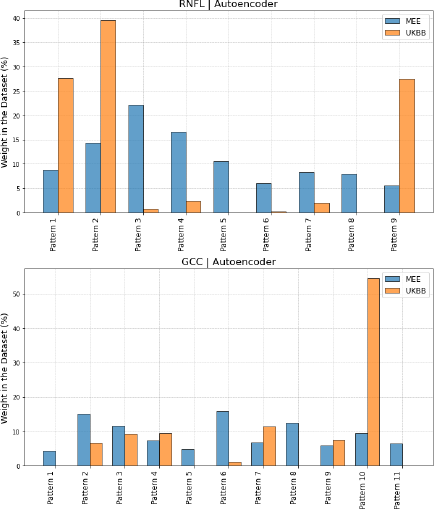
The weight of patterns in the datasets for the autoencoder model

### Encoder Model

As the encoder directly maps the images to feature space, reconstruction errors for the encoder model are not reported.

Similar to the autoencoder model, we applied UMAP on the predictions of the encoder and calculated AIC/BIC scores. The optimal number of clusters was found to be 11 for both GCC and RNFL based on AIC/BIC criteria and the number of clusters with highest correlation (**Figure 7**). The applied GMM on the embeddings is shown in **Supplemental Figure S2**. The labels and representative sample images from each cluster are shown in **Figure 8**. The weight of patterns in each dataset is shown in **Figure 9**. Similar to the results of the autoencoder model, some patterns seen in MEE were not observed when tested in the UKBB dataset (**Figure 9**). For GCC images, patterns 1, 2, 6, 8, 9, and 10 were not observed in the UKBB.

**Fig. 7.**
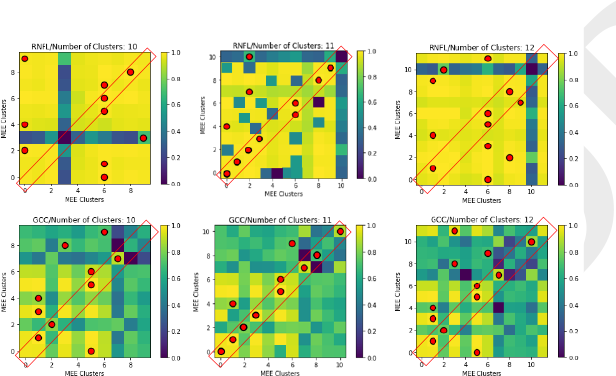
Labels of the data point in the embedding space of GCC images of MOCO model.

**Fig. 8.**
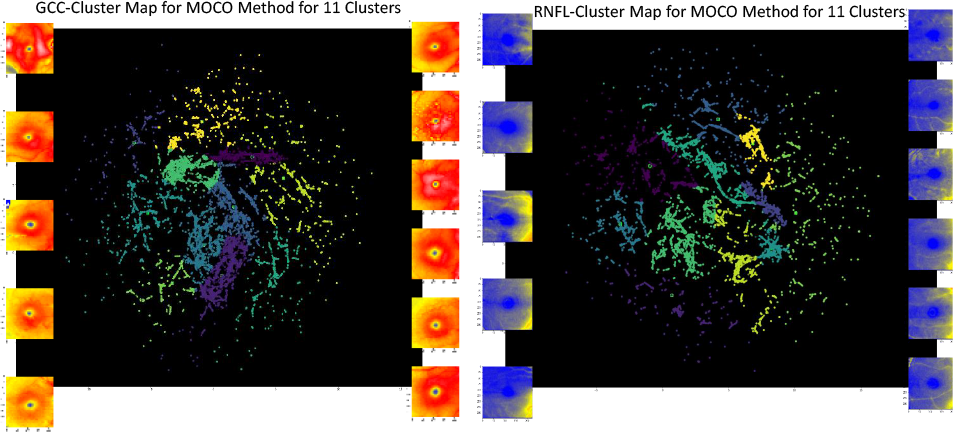
Labels of the data point in the embedding space of GCC and RNFL images of MOCO model.

**Fig. 9.**
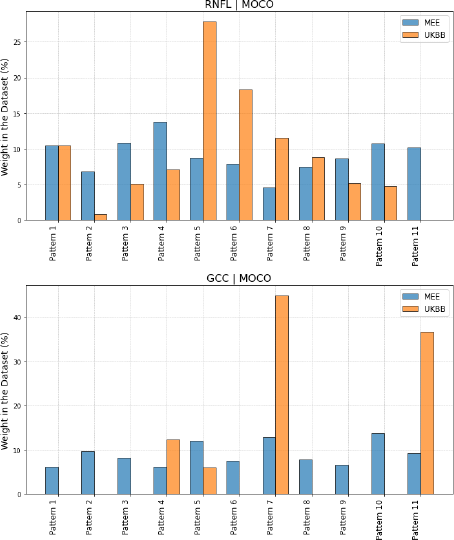
The weight of patterns in the datasets for the encoder model.

### Unsupervised Performance Analysis

We used AIC/BIC scores to identify the optimal number of clusters in our data. Despite the utility of this strategy, we acknowledge its limitations, especially in the context of uncertainty related to the exact number of patterns and the lack of a defined ground truth for evaluating performance. To address these challenges, we introduce a new method for performance assessment that does not require supervision.

To this end, we employed a correlation analysis approach to examine the correlation within and between clusters. This was carried out for images in both pixel and feature space, allowing us to better understand the similarities and differences among images within a single cluster as well as between images in distinct clusters.

We first performed a correlation analysis between images in pixel space. To this aim, we used the normalized crosscorrelation method in which the correlation is calculated based on **Equation** 5:

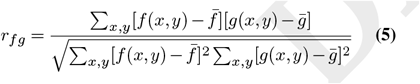

where 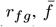 and 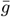 are the correlation, mean values of images *f* and *g*, respectively. however as this method was susceptible to noise, we did not find meaningful results (**Figure 10**). In **Figure** 10a, each cell represents the average correlation between all images of patter *i*: in rows, and pattern *j*: in columns for the RNFL images of MEE dataset for encoder model. As shown, there is no discernible pattern in pixel space correlation. Also **Figure** 10b shows an example of the failure of normalized-cross correlation for correlation analysis; two images that are visually more similar have less correlation (based on normalized-cross correlation) than two images that are less similar visually.

**Fig. 10.**
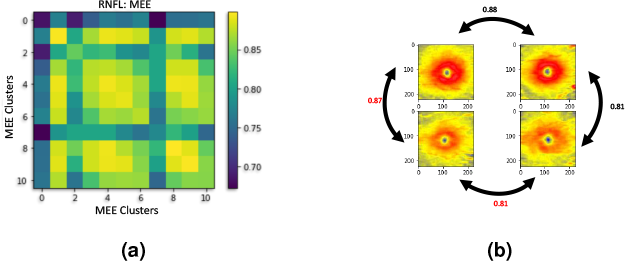
Correlation analysis in pixel space with normalized-cross correlation analysis for RNFL images of the UKBB dataset. Each cell in (a) shows the averaged correlation between the images in each cluster. (b) Example of failure of normalized cross correlation between images. The bottom row shows two similar images from the same pattern that are less correlated (i.e., 0.81) compared to the left column which are images from different patterns (i.e., 0.87). .

We then performed the correlation analysis in the feature space. The correlation analysis in feature space is Pearson correlation (**Equation** 6):

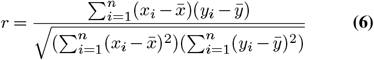

where:

1. *r* is the correlation,
2. *x*_*i*_ and *y*_*i*_ are the i-th elements of variables *X* and *Y*,
3. 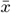 and 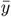 are the means of variables *X* and *Y*,
4. *n* is the number of data points.

The results of the correlation analysis in feature space are shown in **Figures** 11 and 12 . The value for each cell is the average correlation between all images within a cluster and the images in different clusters. Most diagonal cells are brighter than other cells which means the images within a cluster have a higher correlation than the images in different clusters.

**Fig. 11.**
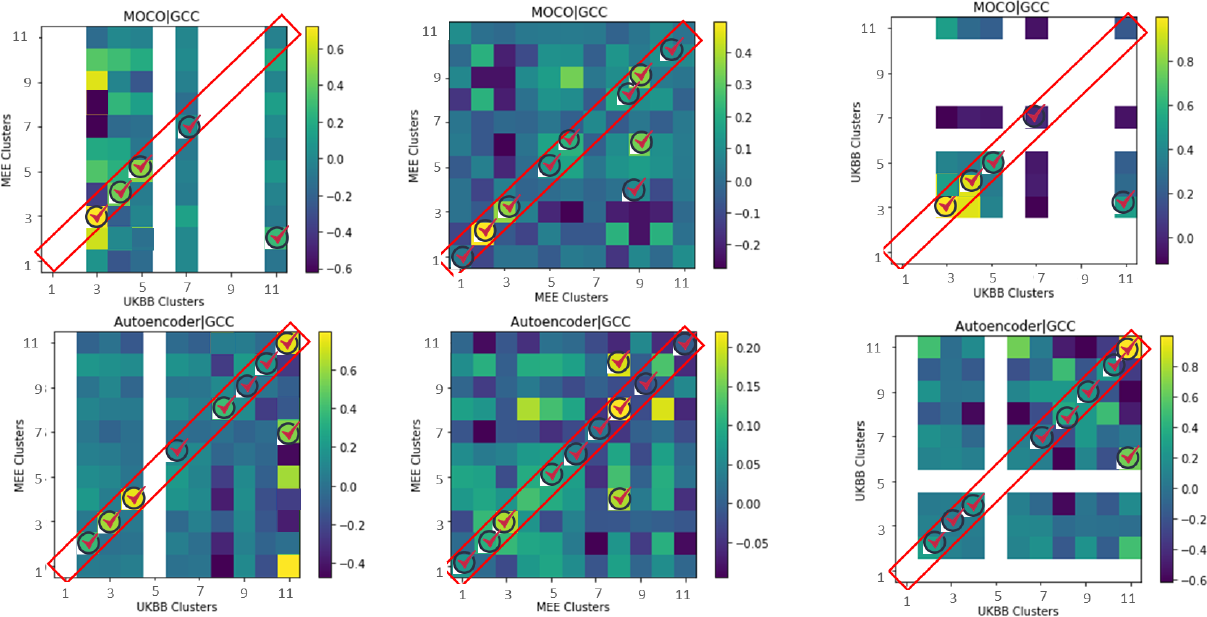
Correlation analysis between the images in feature space with the Pearson correlation analysis for GCC images of MEE and UKBB dataset for both autoencoder and encoder methods.

**Fig. 12.**
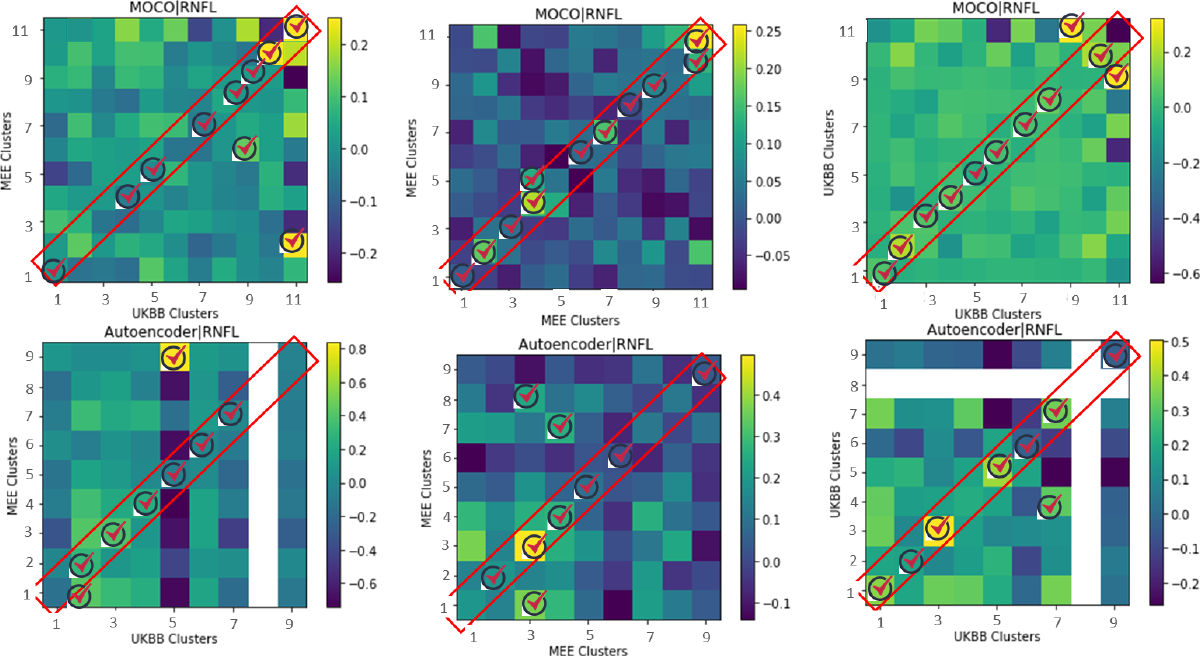
Correlation analysis between the images in feature space with the Pearson correlation analysis for RNFL images of MEE and UKBB dataset for both autoencoder and encoder methods.

### Clinical Associations

To assess the validity of our RNFL and GCC phenotypes, we identified associations between phenotypes and clinical characteristics within the MEE dataset. Correlations between phenotype coefficients and clinical characteristics (IOP, SE, VF MD, CDR, POAG PRS, number of glaucoma medications, and glaucoma severity) can be seen in **Figure 13**.

**Fig. 13.**
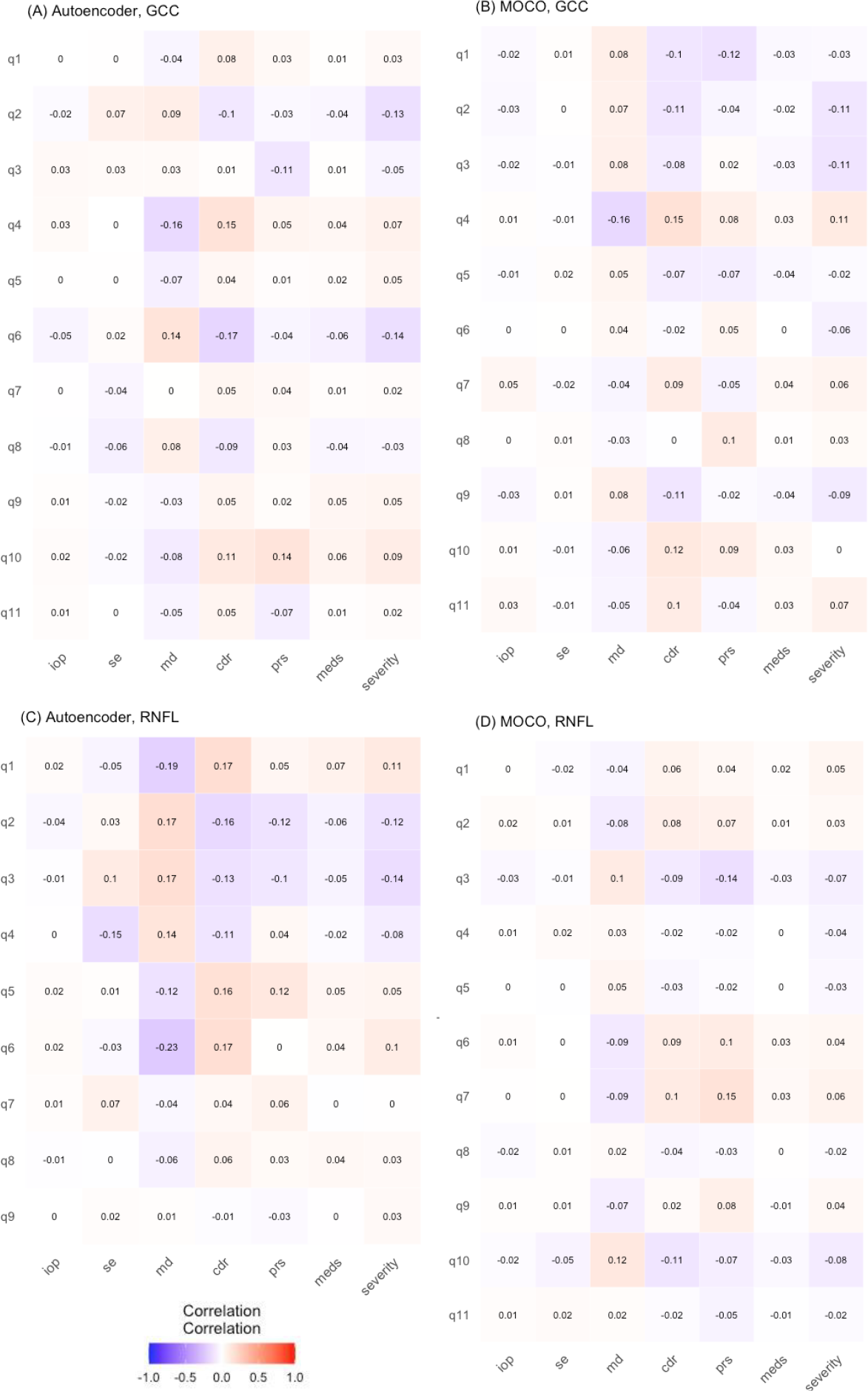
Correlation analysis for MEE phenotypes and clinical characteristics.

For the autoencoder model trained on RNFL thicknesses, phenotype 6 had the highest proportion of severe glaucoma diagnosis codes (ICD 10 diagnosis codes ending in 3 or 4; 49.18%), highest number of average glaucoma medications (1.57 +/- 0.12), highest mean CDR (0.694 +/- 0.237), highest proportion of IOP greater than 24 mmHg (9.99%) and lowest average VF MD (-12.57 +/- 10.1). Phenotype 4 had the highest proportion of highly myopic eyes (SE < -6, 10.73%). For the MOCO model trained on RNFL thickness, phenotype 7 had the highest proportion of severe glaucoma diagnosis codes (41.14%), highest number of average glaucoma medications (1.84 +/- 0.13), highest mean CDR (0.655 +/- 0.220), and lowest average VF MD (-8.278 +/- 7.28). Phenotype 10 had the highest proportion of highly myopic eyes (7.59%). For the autoencoder model trained on GCC thicknesses, phenotype 4 had the highest proportion of severe glaucoma diagnosis codes (39.93%), highest mean CDR (0.654 +/- 0.229), and lowest average VF MD (-9.445 +/- 8.819), while phenotype 9 had the highest number of average glaucoma medications (1.87 +/- 0.11). Phenotype 1 had the highest proportion of eyes with high myopia (7.73%).

For the MOCO model trained on GCC thicknesses, phenotype 4 had the highest proportion of severe glaucoma diagnosis codes (48.42%), highest mean CDR (0.687 +/- 0.221), highest proportion of IOP greater than 24 mmHg (8.41%), lowest average VF MD (-10.95 +/- 9.72) and highest proportion of high myopia (6.85%). Phenotype 7 had the highest number of average glaucoma medications (1.38 +/- 0.11)

## Discussion

Here we demonstrate that unsupervised deep learning can be used to identify imaging phenotypes which have meaningful clinical correlations. We employed a deep learning model, manifold learning, and GMM, and introduced a correlation analysis technique to evaluate the performance of unsupervised phenotyping. Importantly, our new method demonstrated the ability to transfer phenotypes derived from one dataset to another. This work reinforces the potential utility of deep learning models for better understanding of disease pathogenesis and prediction of disease progression.

We demonstrated that our model can be used to identify meaningful patterns of RNFL and GCC thickness amongst patients with glaucoma. In our correlation analysis, we showed that overall, images within one cluster are more correlated than images between different clusters. While pattern detection has been reported based on qualitative analysis in the literature (46) we propose a fully automated pattern recognition in glaucomatous images with a quantitative evaluation method. We also found that some images in clusters had higher correlation with images in other clusters; this may be because GMM assigns labels to each data point based on the highest probability. In some cases, the highest probability pattern may be fairly low (e.g., less than 50%), which means the image may be between multiple clusters. Additionally, GMM assumes that data is distributed in a Gaussian manner, which may not be applicable to real-world datasets such as ours, resulting in imperfect clustering performance. Furthermore, GMM may not perform well with high-dimensional data; as dimensionality increases, the volume of the space increases rapidly such that the available data become sparse making the Gaussian assumption less applicable. Therefore, our model must balance the performance of the GMM and the loss of information due to dataset dimensionality compression.

We compared the use of an autoencoder model and an encoder model based on self-supervised learning; we found that the correlations of the encoder model were higher, in average, compared to those of the autoencoder. This suggests that, while both autoencoders and encoders have demonstrated significant promise in feature extraction tasks, encoders may have an advantage for identifying more meaningful and relevant features. This may be because an encoder focuses on mapping the high-dimensional input data into a lower-dimensional latent space and does not need to reconstruct the original data. This process may result in a more focused, information-rich feature extraction. The encoder concentrates exclusively on data compression, thus preserving more significant patterns in the data and discarding noise. Thus, in certain scenarios, a more simplified or targeted approach like an encoder could outperform more complex models such as autoencoders, further highlighting the importance of problem-specific model selection in machine learning applications.

Importantly, in this work, we demonstrate the ability to successfully transfer OCT phenotypes from one dataset to another with similar reconstruction errors (for autoencoder model) and good correlation of phenotypes (both autoencoder and encoder) in the feature space. Disparate datasets with limited genetic or clinical information pose significant challenges to cross disciplinary and translational research. The ability to transfer phenotypes from one dataset to another will allow for the ability to train and validate phenotypes in a dataset rich with clinical information and apply these phenotypes to a dataset with a genetic repository, thereby potentially increasing the utility of OCT phenotypes to discover novel disease associated genomic loci.

We also demonstrated that our phenotypes have meaningful clinical correlations, including variations in ocular parameters used to assess glaucoma severity such as CDR, visual field mean deviation, and IOP. Specifically, each model (autoencoder vs. MOCO) using different input retinal layers (RNFL vs. GCC) was able to identify a phenotype that correlated with more severe disease, including a more severe ICD code, greater number of glaucoma medications, higher CDR, and lower VF mean deviation. Furthermore, in correlation analysis, several RNFL phenotype coefficients were correlated with SE, a measure for myopia. This may indicate the ability for our models to distinguish RNFL thickness patterns that are affected by myopia; myopia can be associated with “tilting” of the disc due to a longer axial length as is a clinical confounder when assessing patients for glaucoma.

Our study has many notable strengths, including introducing a fully automated pattern recognition using retinal layers obtained from OCT scans, evaluation metrics for unsupervised performance, end to end clinical relevance investigation using the patterns in OCT scans, and transfer capability from one dataset to another. In addition, our MEE dataset size is one of the largest in the U.S.However, retinal thickness layers extracted from OCT images are subject to significant noise, including speckle noise (caused by the interference of multiple scattered light waves), electronic noise (associated with the electronic components of the OCT system, including the detector and analog-to-digital converter), and background noise (due to ambient light and system optics). In order to best correct for noise, we filtered poor quality images and imputed missing pixels. Furthermore, by employing UMAP, we were able to isolate the essential structure of the data and discard aspects of the data that are likely to be noise. UMAP maintains the topological structure of high-dimensional data in lower-dimensional representations, which ensures that the inherent relationships among data points are preserved while the random fluctuations, which often constitute noise, are minimized (47).

Our proposed methodology was effective for identifying clinically meaningful patterns in OCT images using RNFL and GCC thickness maps in patients with glaucoma as a model disease. We were also able to demonstrate the successful transfer of OCT phenotypes from one dataset to another, which may potentially increase the utility of these models for future translational and cross-disciplinary research including better understanding of glaucoma pathogenesis and genetic architecture. Finally, the introduction of a correlation-based assessment represents an exciting development for the performance validation of unsupervised learning methods.

## ACKNOWLEDGEMENTS

This project was supported by the National Institute of Health (NIH) under grants 5K23EY032634-02 and R01EY030575.

## Supplementary Materials

**Table S1.**
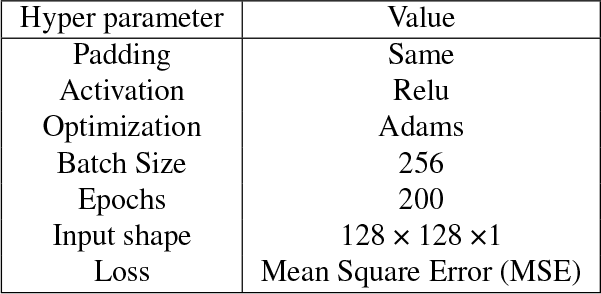
Hyperparameters for training the autoencoder model for feature extraction.

**Table S2.**
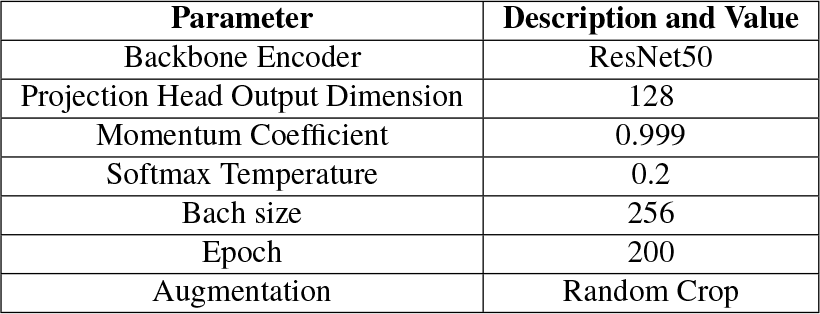
The parameters of Self-supervised learning based on momentum contrast.

| Parameter | Description and Value |
| --- | --- |
| Backbone Encoder | ResNet50 |
| Projection Head Output Dimension | 128 |
| Momentum Coefficient | 0.999 |
| Softmax Temperature | 0.2 |
| Bach size | 256 |
| Epoch | 200 |
| Augmentation | Random Crop |

**Table S3.** UMAP Parameters.

| Parameter | Description or Value |
| --- | --- |
| Number of Reduced Elements | 3 |
| Number of Neighbors | 5 |
| Minimum Distance | 0.0001 |
| Random State | 42 |

**Fig. S1.**
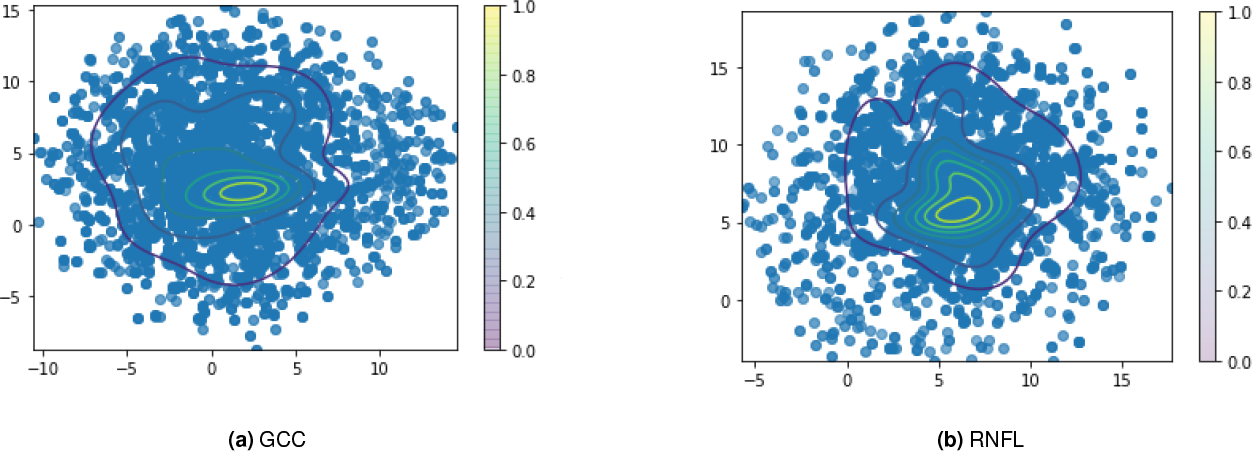
GMM Applied on the Embeddings of GCC and RNFL of MEE dataset for autoencoder model. The plots are in 2D.

**Fig. S2.**
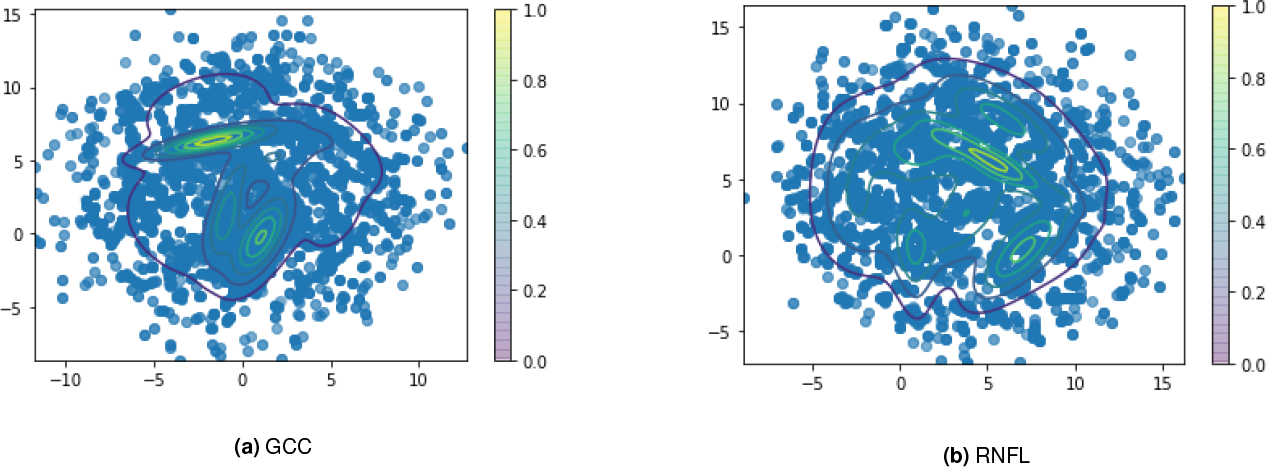
GMM Applied on the Embeddings of GCC and RNFL of MEE dataset for encoder model. The plots are in 2D.

